# DDHD2 is necessary for activity-driven fatty acid fueling of nerve terminal function

**DOI:** 10.1101/2023.12.18.572201

**Authors:** Mukesh Kumar, Yumei Wu, Justin Knapp, Daehun Park, Kallol Gupta, Pietro De Camilli, Timothy A. Ryan

## Abstract

HSP54, a hereditary spastic paraplegia associated with cognitive impairment, is caused by mutations in the neuron-specific triglyceride (TG) lipase DDHD2. Loss of DDHD2 function results in lipid accumulation in human brains^1^ and lipid droplets (LDs) in mouse neurons^2^. In metabolically demanding tissues, TG lipases generate a fatty acid (FA) flux from LDs to fuel mitochondrial ATP production, but neurons are considered unable to use fat as an energy source. Thus, the basis for cognitive impairment driven by DDHD2 loss remains enigmatic. To resolve this paradox, we took advantage of presynaptic sensitivity to metabolic perturbations to determine if FAs derived from LDs could power local β-oxidation to support synaptic functions and whether DDHD2 activity would be required in the process. We demonstrate that nerve terminals are enriched with DDHD2 and blocking its activity leads to presynaptic accumulation of LDs. Moreover, we show that FAs derived from axonal LDs enter mitochondria in an activity-dependent fashion and drive local mitochondrial ATP production allowing nerve terminals to sustain function in the complete absence of glucose. Our data demonstrate that neurons and their nerve terminals can make use of LDs during electrical activity to provide metabolic support when glucose is in short supply.

---

Mammalian brains are metabolically demanding and require a constant supply of combustible fuels to avoid degradation in performance. A decline in glycolytic throughput in the brain is strongly associated with aging and is considered an early predictor of eventual cognitive impairment and neurodegeneration^3-6^. Thus, understanding metabolic pathways the brain uses when glycolysis is limited is likely critical to developing therapeutic approaches to confer resilience during the aging process. The brain, and in particular neurons, have long been considered to rely almost solely on glucose for metabolic support. Other tissues with high metabolic demands, such as muscle, generally show considerable flexibility with respect to the fuel source, and it is paradoxical, considering the brain’s large lipid content and voracious metabolic needs, that it relies on a more restricted choice of oxidizable carbon source for sustaining function. Historic experiments examining whether brain tissue can carry out fat-fueled respiration used tissues that were not physiologically intact^7-9^ and it was incorrectly assumed that acutely provided fatty acids would be as rapidly available for combustion as glucose. The dogma that neurons do not carry out β-oxidation was reinforced by the observation that LDs are rarely seen in healthy neurons. Furthermore, insight into neuronal FA-flux and its metabolic outcome in terms of fueling neuronal functions was limited because most previous studies focused on brain tissues that were a complex consortium of neurons and other cell types^8,10^. The discovery of DDHD2 as a neuron-specific triglyceride lipase warrants a fresh examination of this long-held view since loss of its activity not only results in the buildup of lipid droplets (LDs) within neurons, but also leads to cognitive impairment in both humans^1^ and mice^2^. Here we used various metabolic and molecular sensors to determine that the DDHD2 lipase is highly enriched at synaptic terminals and that electrical activity modulates DDHD2 control of FA-flux from LDs to mitochondria to generate ATP that can support synaptic vesicle recycling in the absence of glucose.

## DDHD2 lipase is enriched in synaptic terminals

Consistent with the fact that genetic deletion of DDHD2 leads to massive accumulation of LDs in mouse neurons, immunocytochemistry with a DDHD2-specific antibody in acute brain slices (Fig. 1a) shows that this protein is present throughout the hippocampus (Fig. 1b). The staining was evident in areas enriched with large synaptic terminals such as hilus, as visualized with a synapsin antibody (Fig. 1c). Quantification revealed that the fluorescence signal of DDHD2 and synapsin in the hippocampal hilus overlapped with a Pearson’s colocalization coefficient of 0.62 ± 0.016 (Fig.1f). To further characterize the subcellular distribution of DDHD2 and determine its distribution in axons, we carried out immunocytochemistry analysis in sparsely plated dissociated primary rat hippocampal neurons which showed that in axons, DDHD2 accumulates to the same extent as synapsin in nerve terminals (Fig. 1d, f, g). Neurons in which DDHD2 was ablated (by small hairpin RNA [shRNA] expression) showed much lower levels of staining (Supp. Fig. 1a, b) confirming the specificity of the antibody. Additionally, co-expression of GFP-DDHD2 and mRuby-synapsin showed strong co-localization in hippocampal neurons, with a Pearson’s coefficient 0.60 ± 0.048 (Fig 1e, f). Thus, we conclude that DDHD2 is distributed throughout neurons, but it in axons is highly enriched in nerve terminals.

**Figure 1.**
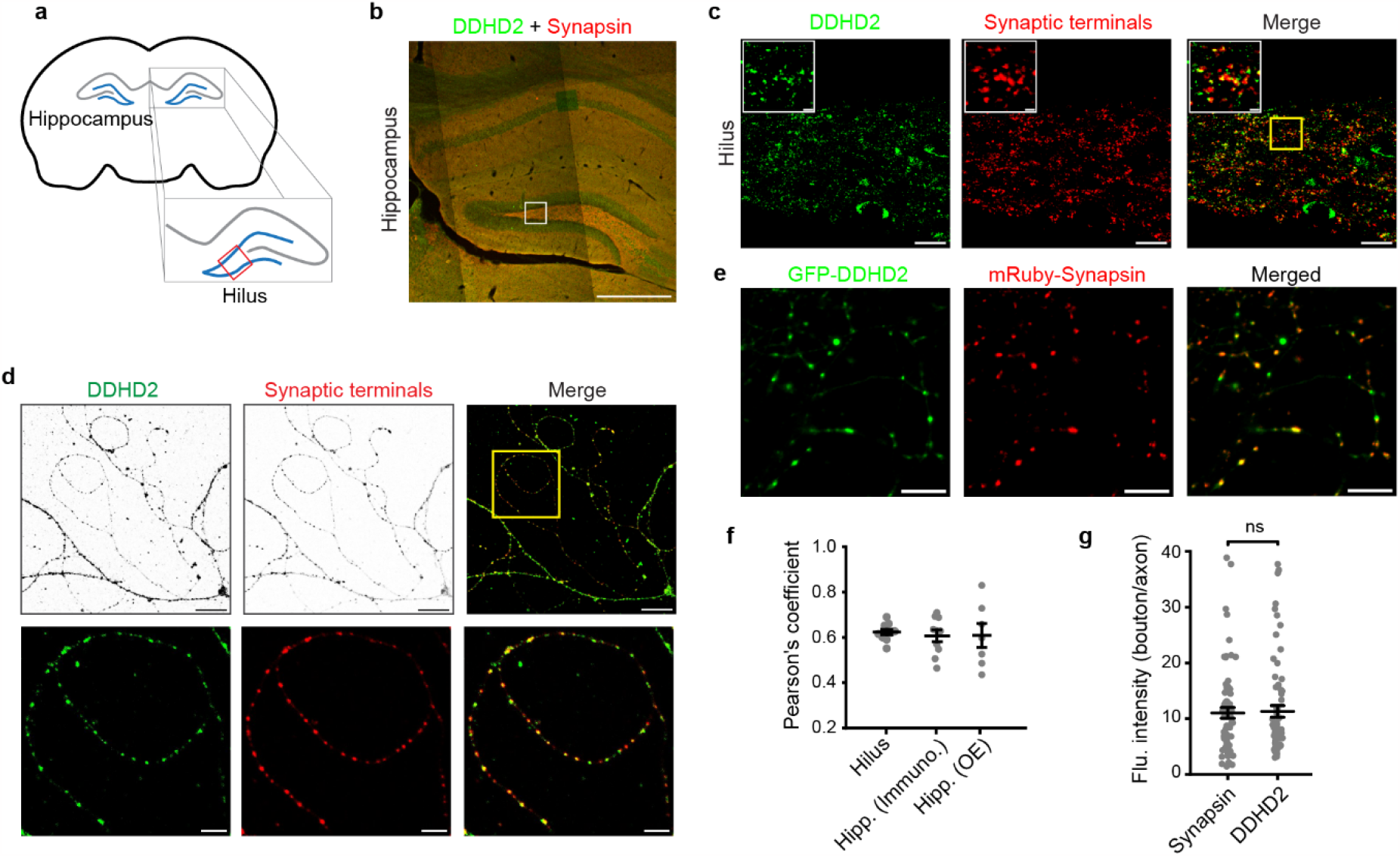
DDHD2 lipase at the synaptic terminals. **a**, Schematic of coronal section of a rodent brain. Hilus is boxed in the zoomed-in part of the hippocampus. **b**, Confocal micrograph of a hippocampus derived from adult (28 day old) mouse brain, immunolabeled with DDHD2 and synapsin (synaptic terminal) antibodies. scale bar: 500 μm. **c**, Confocal micrographs of the zoomed-in region of hilus as marked in (b), immunolabeled with DDHD2 and synapsin antibodies, scale bar: 20 μm. 15 μm x 15 μm area (yellow box in the right panel) of hilus are shown in insets. scale bar: 2 μm. **d**, Confocal micrographs of dissociated hippocampal neurons immunolabeled with DDHD2 and synapsin antibodies. scale bar: 20 μm. Lower panel shows 50 μm x 50 μm area (yellow box in upper right panel). scale bar: 5 μm. **e**, Confocal micrographs of dissociated hippocampal neurons expressing GFP-DDHD2 and mRuby-Synapsin. scale bar: 10 μm. **f**, Colocalization of immunolabeled DDHD2 and synapsin in hilus (15 μm x 15 μm, n=10), hippocampal neurons (65 μm x 65 μm, n=10) and DDHD2 & synapsin overexpressing hippocampal neurons (135 μm x 135 μm, n=7) as quantified by Pearson’s coefficient. Data is represented as mean ± SE. g, Relative enrichment of synapsin and DDHD2 at the synaptic terminals compared to adjacent axonal processes of hippocampal neurons in (d). Data is represented as mean ± SE and *p*-value (ns: not significant) was determined using unpaired samples t-test.

### Inhibition of DDHD2 activity causes accumulation of LDs at synapses

Mice treated with KLH45 (a selective DDHD2 inhibitor) over a period of 4 days, show significant elevation of brain TGs, similar to changes observed upon genetic ablation of this enzyme^2^. We treated sparsely plated dissociated hippocampal neurons with KLH45 for 24 hrs, and immuno-labeled synaptic terminals with a synapsin antibody and the neutral lipid stain BODIPY(493/503), a convenient marker of LDs^11^ (Fig. 2a, b). These experiments revealed that ∼70 % of synaptic terminals were positive for BODIPY (Fig. 2c), demonstrating that most nerve terminals appear to be constantly consuming LD TGs. Electron microscopy-based ultrastructural analysis confirmed the presence of electron-dense LDs close to synaptic vesicle clusters (Fig. 2d). We further verified that LDs typically appear at nerve terminals with resident mitochondria, by triple staining of KLH45 treated neurons with BODIPY (LDs), anti-TOMM20 (mitochondria) and anti-synapsin (synaptic terminals) antibodies (Fig. 2e, f). The line intensity profile of 50-100 μm stretch of axonal processes shows frequent peaks of BODIPY and TOMM20 at synaptic terminals (Supp. Fig. 2a, b).

**Figure 2.**
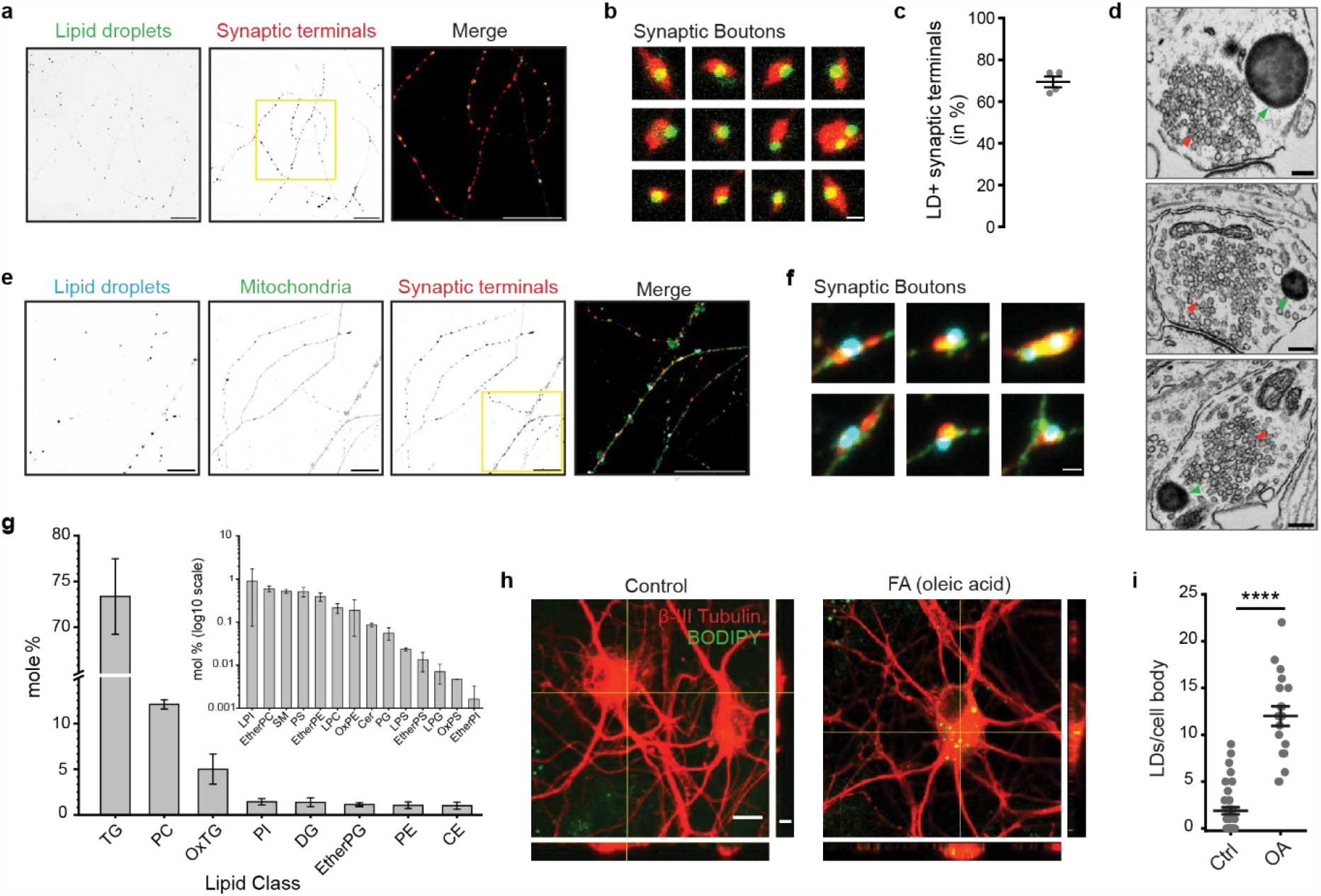
Inhibition of DDHD2 leads to LD accumulation at synaptic terminals. **a**, Confocal micrograph of the KLH45-treated hippocampal neurons immunolabeled with synapsin (synaptic terminal) antibody and stained with BODIPY (LDs) dye. Yellow boxed region is shown as a merged zoom-in image. scale bar: 20 μm. **b**, Zoomed-in region of 12 individual synaptic boutons from (a). scale bar: 1 μm. c, Quantification of synaptic terminals detected positive for BODIPY stained LDs. Data is represented as mean ± SE, N=4. **d**, Electron micrographs of KLH45-treated (18 DIV) hippocampal neurons. LDs and synaptic vesicles are indicated by green and red arrowheads respectively. scale bar: 200 nm. **e**, Confocal micrographs of KLH45-treated hippocampal neurons immunolabeled with TOMM20 (mitochondria) and synapsin (synaptic terminals) antibodies, and stained with MDH/AUTODOT (LDs) dye. Yellow boxed region is shown as a merged zoom-in image. scale bar: 20 μm. **f**, Zoomed-in region of 6 synaptic boutons from (e), scale bar: 1 μm. **g**. Lipids extracted from the isolated LDs of cortical neurons were detected and quantified (in mole %) using LC-MS. Minor lipids are shown (note the logrithmic scale) in the inset. Data is represented as mean ± SD. **h**, Confocal micrographs of control and oleic-acid fed hippocampal neurons stained with anti β-III tubulin antibody (neuron) and BODIPY (LD), scale bar: 10 μm. Orthogonal sections are shown to indicate globular LDs in the cell body, scale bar: 5 μm. **i**, Quantification of the number of LDs in the cell body of control and oleic acid fed hippocampal neurons. Data is represented as mean ± SE and *p*-value (**** ≤ 0.0001) was determined using unpaired samples t-test.

We carried out a biochemical purification of these LDs in our neuronal cultures using a sucrose density gradient (Supp. Fig. 2c) to further characterize their lipid content. As expected, the outer surface of the purified LDs stained positively with the membrane dye CellMask^12^, while the core stained positively for BODIPY (Supp. Fig. 2d, e) and had an average diameter of 1.2 μm (Supp. Fig. 2f). Quantitative LC-MS of these LDs showed that they consisted of 73.35 ± 4.12 % TGs, 1.01 ± 0.39 % CE, and 12.09 ± 0.5 % PC (of mol respectively; Fig. 2g; Supp Fig. 2g). We also identified a lower abundance of other lipids classes (Fig. 2g, Supp. Fig. 2g, h). A detailed method for purification and lipidomic profile of the neuronal LDs can be accessed elsewhere^13^.

This data demonstrates that under steady-state metabolic conditions, TGs are likely continuously being synthesized and consumed, as blocking DDHD2 activity shifts the balance to a net accumulation of LDs. We tested an alternative approach to tilt the balance towards LD accumulation, which was to provide excess FAs to the growth medium. BODIPY staining carried out to neurons whose feeding media was supplemented with excess oleic acid for 24 hours showed that altering the amount of available FAs also leads to a net accumulation of LDs in neurons (Fig. 2h, i).

### Metabolic demand in neurons triggers triglyceride mobilization from LDs

In other tissues, TGs are stored in LDs to serve as an alternative fuel source when glucose availability is limited. Lipases are necessary for TG hydrolysis, converting them into FAs in the cytosol. Subsequently, these FA molecules are converted into fatty acyl-CoA and delivered to the mitochondrial matrix via the mitochondrial carnitine palmitoyl-transferases (CPTs) for oxidation. To determine if FAs derived from LDs in axons are delivered to axonal mitochondria, we used a pulse-chase strategy with fatty acid covalently conjugated with BODIPY 558/568, Red-C12 (Fig. 3a). A 24-hour pulse of Red-C12 was delivered to dissociated hippocampal neurons in the presence of KLH45 to allow it to accumulate in LDs (Supp. Fig. 3a-c) followed by removal of KLH45 during chase period. The amount of Red-C12 accumulated in mitochondria was assessed by measuring the fluorescence intensity of Red-C12 that overlapped with MitoTracker Green^14^. 24 hours after pulsing in Red-C12 during DDHD2 block, Red-C12 showed both diffuse axonal labeling as well as a punctate distribution that were positive for LD markers (Supp. Fig. 3a), with minimal co-localization with mitochondria (Fig. 3b, see 0 hr. of Fig. 3c), however after removing the DDHD2 inhibitor, we observed a gradual transfer of Red-C12 into mitochondria during the chase period (Fig. 3b, see 1.5-4.5 hr. of Fig. 3c). In contrast, including the CPT1 inhibitor etomoxir during the chase period prevented transfer of Red-C12 to mitochondria (Fig. 3b, c).

**Figure 3.**
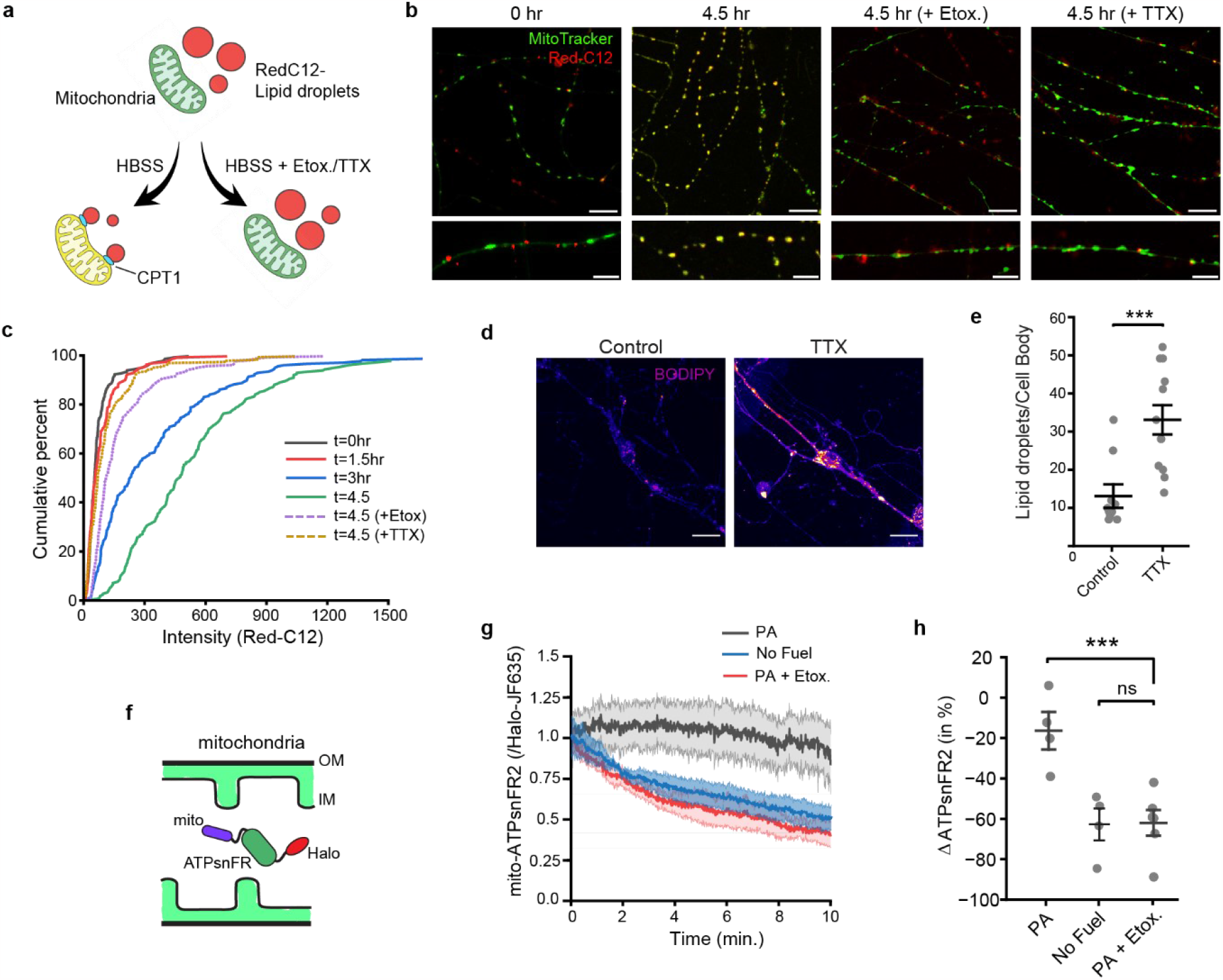
Synaptic activity and CPT1 facilitates FA transfer to mitochondrial matrix to maintain ATP level in absence of glucose. **a**, Schematic to show the accumulation of Red-C12 in LDs after treatment with KLH45. CPT1 & 2 (CPT2 not shown in schematic) on mitochondrial membranes facilitates transfer of Red-C12 to mitochondrial matrix in HBSS media. **b**, Confocal micrographs show transfer of Red-C12 to mitochondria within 4.5 hrs. in absence of external glucose. Presence of Etomoxir or TTX in glucose-free HBSS media inhibits the transfer of Red-C12 to mitochondrial matrix. scale bars: 10 μm (upper panels) & 5 μm (lower panels). **c**, Cumulative percent transfer of Red-C12 to the mitochondrial matrix at 0 hr., 1.5 hr., 3 hrs., and 4.5 hrs. after HBSS media change. Dotted lines represent reduced Red-C12 transfer to mitochondrial matrix when etomoxir or TTX was present in HBSS. **d**, Confocal micrographs of control and TTX (72 hrs.) treated hippocampal neurons stained with BODIPY. scale bar: 20 μm. **e**, Quantification of LD number in cell body of control and TTX treated neurons. Data is represented as mean ± SE. *p*-value (*** ≤ 0.001)) was determined by unpaired samples t-test. **f**, Schematic illustration of C-terminally Halo-tagged ATP sensor (ATPsnFR2) targeted into mitochondrial matrix with the help of four repeats of N-terminal leader sequence, mito. **g**, ATP levels (normalized using JF635) in the mito-ATPsnFR2-Halo expressing hippocampal neurons in absence of any fuel, PA induced or CPT1 inhibited condition (by Etomoxir) to the PA induced neurons. Data is represented as mean ± SE for N=4 (each condition). **h**, Relative change (in %) in the ATP levels after 10 minutes of perfusion with the media as described in (g). Data from N=4 for each condition is represented as mean ± SE, *p*-value (** ≤ 0.001 for PA and PA + Etox.) was determined using unpaired samples t-test. ns signifies non-significant change in mean values.

### Electrical activity controls LD accumulation by modulating the transfer of FAs from LDs into mitochondria

We previously showed that electrical activity is tightly coupled to ATP synthesis in neuronal axons to meet the demands of nerve terminal function^15^ through both upregulation of glucose uptake^16^ and mitochondrial respiration^17^. To determine if electrical activity might impact LD storage or usage, we treated dissociated neuronal cultures with the Na^+^ channel blocker TTX for 72 hours, and examined the LD content compared to untreated neurons using BODIPY (Fig 3d). These experiments demonstrated that blocking chronic electrical activity leads to a significant accumulation of LDs both in cell somas and in neuronal processes (Fig. 3d, e). This result implies that electrical activity modulates some combination of LD formation or consumption. To see if LD consumption might be modulated, we used our Red-C12 pulse-chase approach but blocked electrical activity during the chase period with TTX (after removal of the DDHD2 inhibitor). These experiments revealed that blocking electrical activity suppressed the transfer of FA into the mitochondrial matrix (Fig. 3b, c), demonstrating that consumption of LDs at nerve terminals is modulated by electrical activity. These experiments demonstrate that the abundance of LDs in neurons is determined by both the availability of FAs, as well as electrical activity which in turn modulates the consumption of LDs.

### Fatty acids derived from axonal LDs drive β-oxidation and axonal mitochondrial ATP production

We made use of a newly developed large dynamic range genetically encoded ATP sensor targeted to the mitochondrial matrix, mito-iATPSnFR2^SP54, 18^, expressed in primary neurons to examine ATP production in the mitochondrial matrix (Fig. 3f) under different fuel conditions. Following acute removal of glucose, axonal mitochondrial ATP declines slowly for the next 10 minutes, as expected since pyruvate production via glycolysis should be largely absent. However, in neurons that had been pre-fed with palmitic acid for 4 hours mitochondrial ATP levels were sustained over the same period in the complete absence of glucose. In contrast if the CPT1 inhibitor, etomoxir was applied at the same time as removing glucose, the decline in mitochondrial ATP level in palmitic acid pre-fed neurons was identical to the control neurons without any fuel supplement (Fig. 3g, h). Similar results were obtained in neurons where LDs were induced by blocking DDHD2 activity with KLH45 for 24 hours. When glucose and KLH45 are removed, ATP levels are sustained, but including etomoxir under these conditions lead to ATP decline over a 10-minute period (Supp. Fig. 3d, e). These experiments demonstrate that axonal mitochondria can sustain mitochondrial ATP production using FAs derived from LDs in the complete absence of glucose.

### LDs support synaptic vesicle recycling in hypoglycemic conditions

Our data demonstrate that axonal DDHD2 lipase activity facilitates the release of FAs from LDs, fueling mitochondrial ATP production. A critical question is whether the ATP produced from LD derived FAs can support synaptic function. To test this, we used endocytic retrieval of vGlut1-pHluorin at synaptic terminals as a measure of metabolic endurance. In the absence of an oxidizable carbon source, we previously showed that pHluorin-tagged SV proteins fail to recycle, instead accumulating on the plasma membrane or in an alkaline compartment^15,16^. Here we deployed a more stringent synaptic endurance assay, consisting of repeated 50 action potential bursts of firing at minute intervals (Fig. 4a). Under sufficient fuel conditions, nerve terminals can sustain synaptic vesicle (SV) recycling through at least five such rounds of stimulation, however when glucose is removed vesicle recycling fails within the first 2-5 rounds. (Supp. Fig. 4a). In contrast, when LDs were allowed to accumulate TGs by transiently blocking DDHD2 with KLH45, or by pre-feeding with PA for 4 hours, SV recycling could now be sustained in the complete absence of extracellular glucose (Fig. 4b, Supp. Fig. 4a). Adding KLH45 acutely had no impact on SV recycling in glucose, but in neuron pre-fed PA, SV recycling arrested within 1-2 rounds of stimulation (Fig. 4c). As would be expected from a fuel that requires the electron transport chain, incubation in the with F_0_-F_1_ ATPase inhibitor (oligomycin) completely arrests SV recycling that depends on LDs (Supp. Fig. 4b). These data indicate that pre-feeding neurons with FAs fuels ATP production via lipolysis of LD resident TGs. Consistent with the need to assemble LDs, brief (5 min.) incubation of neurons with extracellular FAs fail to sustain SV recycling in the absence of glucose (Supp. Fig. 4c). Moreover, inhibition of CPT1 to FA-fed neurons by etomoxir acutely inhibited SV recycling (Fig. 4d). To rule out the possibility that the improvement may have been caused by metabolites arising from glial cells in the same dish, LD-driven SV recycling fails in when the entire dish of cells (neurons & glia) was previously infected with a lentivirus, driving expression of an shRNA targeting CPT2 but was restored when an shRNA-insensitive variant was re-expressed in neurons (Supp. Fig. 4d-h). These data demonstrate that presynaptic function can be sustained purely from axonal LD-derived FAs.

**Figure 4.**
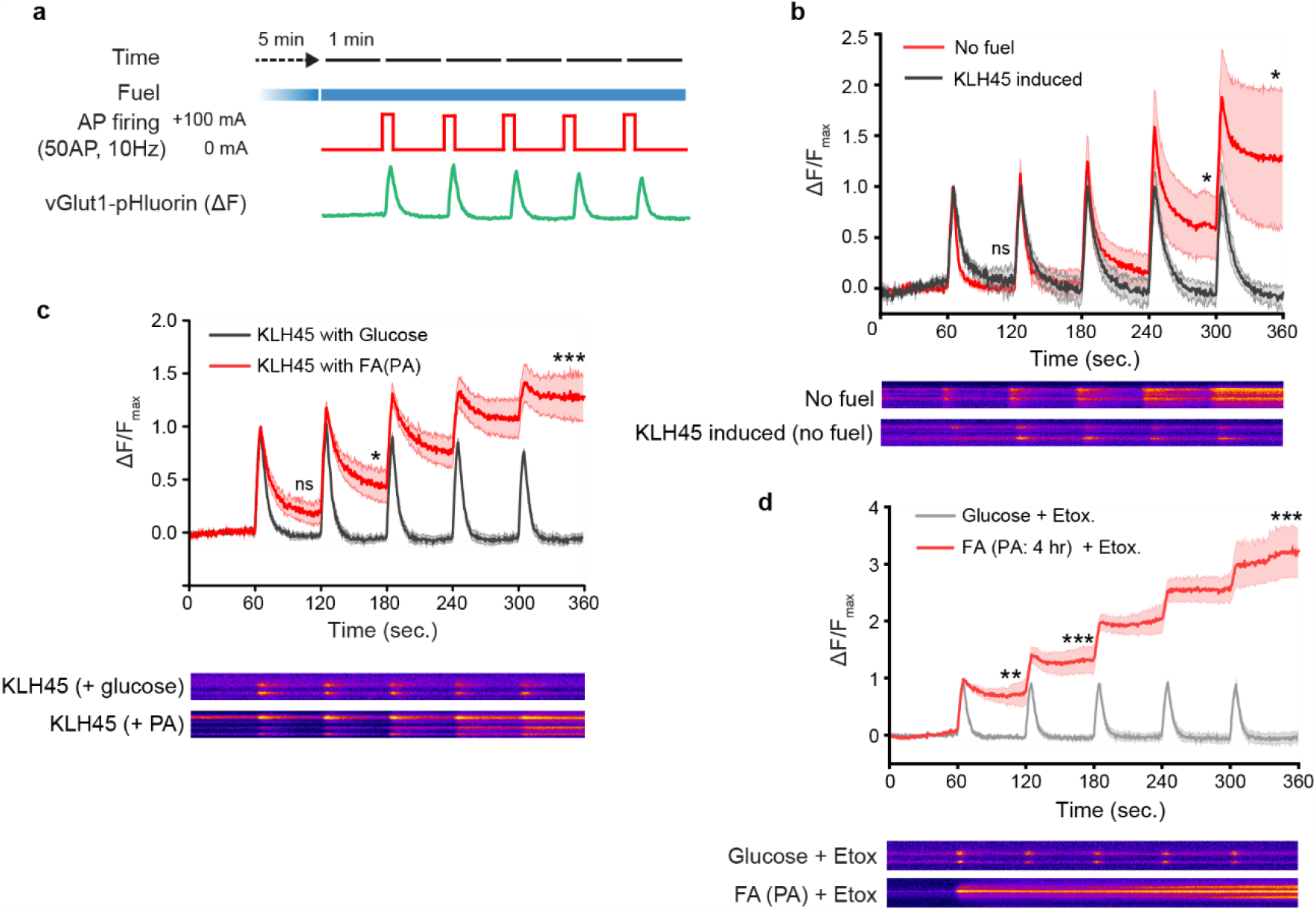
Lipid droplet induced neurons show improved synaptic endurance. **a**, Experimental timeline. vGlut1-pHluorin expressing dissociated hippocampal neurons were perfused with desired media for five minutes (or indicated otherwise) followed by action potential (AP) firing (+100 mA, 50 AP, 10 Hz) at 60 second intervals. The fluorescence intensity shift of pHluorin (ΔF) at the synaptic boutons (ROIs: 12-18 μm^2^) due to exo-and endocytosis of SVs were recorded and normalized with the peak after completion of first AP firing. **b**, Fluorescence intensities at synaptic boutons after periodic AP firing in absence of any external fuel (labeled No fuel) and KLH45 induced hippocampal neurons. Unlike no fuel condition, TG loaded neurons sustain 5 rounds of SV recycling. Data is represented as mean ± SE, *p*-value (* ≤0.05) was determined using unpaired samples t-test. **c**, Fluorescence intensities at synaptic boutons after periodic AP firing in KLH45 with glucose or PA fed hippocampal neurons. Data is represented as mean ± SE, *p*-values (* ≤0.05, *** ≤0.001) were determined using unpaired samples t-test. **d**, Fluorescence intensities at synaptic boutons after periodic AP firing in glucose or PA fed hippocampal neurons in presence of Etomoxir. Data is represented as mean ± SE, *p*-values (** ≤0.01, *** ≤0.001) were determined using unpaired samples t-test. ns in ‘b’ and ‘c’ signifies non-significant changes in mean values. Representative kymographs for the changes in the vGlut1-pHluorin intensities at the synaptic boutons are shown at the bottoms of their respective traces in b, c, and d.

## Discussion

The discovery that loss of DDHD2 function leads to a profound accumulation of LDs in neurons throughout the brain overturns a long-held view that neurons do not store, and therefore do not use TGs, as part of their metabolic program. Since loss of DDHD2 function in humans and mice leads to significant cognitive impairment, these results imply that this branch of metabolic support serves a critical role in sustaining neuron health and function, but previous studies did not examine what function these LDs normally consumed by DDHD2 play. Here we demonstrate conclusively that although at steady-state LD abundance in neurons is very low, in agreement with most historic observations, that this is the result of a constant flux of FAs through the LD pathway. Remarkably we find that this pathway plays a prominent role in axons, as DDHD2 appears highly enriched in nerve terminals and inhibiting this enzyme leads to the appearance of LDs in the majority of nerve terminals. We show that this flux itself is controlled by neuronal electrical activity and that FAs once liberated from TGs in LDs are transferred into the mitochondrial matrix, which then support mitochondria-based ATP production.

Our work now brings into focus several key questions that are likely central to brain bioenergetics, in particular the source of FAs that are normally used to build LDs and subsequently consumed by DDHD2 and the route for delivery of FAs into the neuron cytoplasm. One class of attractive candidates that might act as the extracellular conduit for FAs are lipoprotein particles (LPs), a common vehicle for shuttling sterols and TGs throughout the body. Astrocytes are well known to produce LPs that are secreted into the extracellular space in the brain^19^. One of the main protein adaptors on astrocyte-derived lipoprotein particles is ApoE, the allelic variation of which is known to significantly impact Alzheimer’s disease risk^20^. Efficient extraction of TGs from lipoprotein particles generally requires that the receiving cell express lipoprotein lipase (LPL), a secreted protein, that normally acts at the surface of a cell to extract FAs for delivery to surface FA transporters. Notably most neurons in the brain express LPL, and genetic deletion of neuronal LPLs leads to a spectrum of neuronal dysfunction ^21,22^. It has long been known that in muscle, FAs and glucose can co-regulate each other’s use as a fuel, such that one fuel is preferentially used and inhibits the combustion of the other, known as the Randle Cycle ^23,24^. It is interesting to consider that such cross-regulation might be operational in neurons as well and it will be interesting in the future to understand additionally how this potential cross-regulation might be in turn be impacted by electrical activity. In the future it will be interesting to determine the cell biological routes of how TG-derived FAs are transferred to the surface of mitochondria, and the underlying machineries that support this as well as to determine the suitability of LPs as an extracellular route of TG delivery to neurons. It is interesting to consider that the ability of neurons to make use of LD-derived FAs for local presynaptic ATP production may become increasingly important in aging brains as the access to glucose in the brain might become more limited^25^.

## Methods

### Animals

Wildtype strains of Sprague Dawley rats (Charles River code 400, RRID: RGD_734476) and mouse were used to culture neurons. All the animal experiments were performed in accordance with the protocol reviewed and approved by the Institutional Animal Care and Use Committee (IACUC) of Weill Cornell Medicine or Rockefeller University.

### Chemicals

All the chemicals used in the study were of analytical grade and procured from Sigma Aldrich unless specified otherwise in the methods. The solutions and buffers were prepared with deionized water of resistivity 18.2 MΩcm^-1^ or above. The pharmacological inhibitors were dissolved in Dimethyl sulfoxide, DMSo (Sigma, D2650).

### Primary cultured neurons and transfection

Glass coverslips were coated overnight with 10 μg/ml solution of poly-ornithine (Sigma-Aldrich, P-3655). The hippocampal regions (CA1-3) were isolated from neonatal rat pups aged 0-1 day, without gender-based distinction. The dissociated hippocampal neurons were cultured in MEM (ThermoFisher Scientific, 51200038) containing 29 mM glucose, 0.1 mg/ml bovine transferrin (Millipore, 616420), 24 μg/ml insulin (Millipore Sigma, I6634), 1% Glutamax (Thermo Fisher, 35050061), 5% fetal bovine serum (Atlanta Biologicals, S11510), 2% N-21 (R&D Systems, AR008), and 4 μM cytosine β-D-arabinofuranoside (Millipore Sigma, C6645). The cultures were maintained at 37°C in a humidified incubator with 95% air and 5% CO_2_ for the next 21 days until used for the experiments. The neurons were transfected using calcium phosphate method after 7 days of plating. Imaging was performed between 7 to 14 days post-transfection. The detailed stepwise protocols for hippocampal culture, transfection, and imaging conditions are described elsewhere (dx.doi.org/10.17504/protocols.io.ewov1qxr2gr2/v1).

### IHC with brain sections

Brain of adult (28-30 day) mouse was dissected and submerged in a 4% (w/v) PFA (Thermo Scientific, J19943K2) and 15% (w/v) sucrose solution overnight at 4 °C for fixation. Subsequently, the brain was mounted on cryostat base with sucrose (Sigma, S0389), and 30 μm thick coronal section spanning CA1-CA3 were prepared. The sections were washed 3 times (5 minutes each) with PBS to remove the cryo-protectant. The antigens were retrieved by heating the sections at 95 °C in citric acid buffer (10 mM citrate, 0.05% tween-20, pH 6) for 20 minutes. The retrieved sections were transferred to PBST (0.5% tween-20 and 0.1 % triton X-100 in PBS). The washing step with PBST was repeated 3 times (10 minutes each) on shaker at room temperature. The sections were then submerged in blocking buffer (0.1% BSA, 2% NDS, 0.1% Triton X-100, 0.05% tween-20 and 50 mM glycine in PBS) for 1 hour at room temperature. The blocking buffer was exchanged with ASE (antibody signal enhancer) solution (10 mM glycine, 0.1% of 30% H_2_O_2_, 0.05% tween-20, 0.1% Triton X-100 in PBS) containing the recommended concentration of primary antibodies. The sections were incubated in the primary antibodies overnight at 4°C in a humidified chamber. The sections were washed 3 times (10 minutes each) with PBST followed by incubation in recommended concentration of labelled secondary antibodies prepared in PBST for 1 hour at room temperature in a humidified chamber. The secondary antibody was aspirated off and the sections were again washed with PBST for 3 cycles (10 minutes each). The excess washing buffer was soaked off with the corner of a tissue paper and ∼100 μl of mounting media was immediately dispensed over the stained brain sections. A clean refractive index-matched coverslip was carefully placed on the sections and edges were mounted with clear nail-polish.

### Immunocytochemistry with primary hippocampal neurons

Hippocampal neurons were sparsely cultured from post-natal day 0-1 pups on glass coverslips coated with poly-d-lysin (Sigma-Aldrich, P7886). When neurons reached 14 days *in-vitro* (DIV-14), they were fixed using a solution of 4% PFA and 10% sucrose in PBS for 10 minutes at room temperature. The fixed neurons were rinsed 3 times with PBS (5 minutes each). The fixed neurons were permeabilized with 0.25% Triton X-100 in PBS for 5 minutes, followed by 3 cycles of PBS washes (5 minutes each). The permeabilized neurons were blocked with 5% goat serum (Sigma-Aldrich, G9023) in PBS for 1 hour at room temperature. The blocked neurons were gently washed once with PBS for 10 min. Primary antibodies were diluted (following manufacturer’s instructions) in 5% goat serum prepared in 0.05% Triton X-100 containing PBS, applied to blocked neurons, and incubated overnight at 4°C in a humidified chamber. The following day, the neurons were washed 3 times with PBS (5 minutes each). Subsequently, secondary antibodies (conjugated to Alexa 488 or Alexa 546, Invitrogen) were applied to the neurons typically in a 1:500 dilution prepared in a solution of 0.05% Triton X-100 in PBS and incubated for 1 hour at room temperature in a humidified chamber. Further, the neurons were subjected to two rounds of PBS washing, each for 5 minutes. To stain the lipid droplets, the third wash was performed for 10 minutes in PBS containing 1 μg/ml BODIPY (Invitrogen, D3922) or 100 μM MDH (Abgent, SM1000a) followed by a gentle wash with PBS for 10 minutes. Finally, the glass coverslips containing stained neurons were mounted on glass slides using mounting media that contained BODIPY or MDH to compensate any potential bleaching during microscopy.

### Electron microscopy with primary cultured neurons

The hippocampal neurons were dissociated and plated densely confined within 5 mm cylinder mounted on poly-ornithine coated glass coverslips. The neurons were fixed for 1 hour at room temperature using a solution containing 2.5% Glutaraldehyde and 2 mM CaCl_2_ in 0.1 M sodium cacodylate buffer. The fixed neurons were washed 4 times each for 5 minutes with 0.1 M sodium cacodylate buffer containing 2 mM CaCl_2_. Subsequently, the neurons were post-fixed in 2% OsO_4_ and 1.5% K_4_Fe(CN)_6_ + 2mM CaCl_2_ in 0.1 M sodium cacodylate buffer. The samples were en bloc stained with 2% aqueous uranyl acetate and 2 mM CaCl_2_ (J.T.Baker), dehydrated, and embedded in Embed 812. Ultrathin sections 50-60 nm thickness were observed in a Talos L 120C TEM microscope at 80 kV. Final images were captured using Velox software and a 4k × 4K Ceta CMOS Camera (ThermoFisher Scientific). Unless otherwise specified, all reagents for electron microscopy were acquired from EMS (Electron Microscopy Sciences), Hatfield, PA.

### Lipidomic analysis of isolated neuronal lipid droplets

Lipid droplets were induced in the dissociated cortical neurons and purified using a sucrose density gradient. The detailed method for induction, purification and biophysical analysis of the neuronal lipid droplets is published elsewhere^13^ (https://doi.org/10.1101/2023.12.13.571527). The LD samples were subjected to MTBE extraction with spiked Avanti Splashmix as the internal standards. The organic phase was collected, dried, and subsequently resuspended in 8-Butanol. 3 μl of this mixture was loaded onto an Agilent Poroshell 120 (EC-C18 2.7 μm, 1000 bar, 2.1 x 100 mM) column using an Agilent 1290 Infiniti II LC. Mobile Phase A (60% Acetonitrile, 40% H_2_O, 7.5 mM Ammonium Acetate) and mobile phase B (90% IPA, 10% Acetonitrile, 7.5 mM Ammonium Acetate) were used to separate peaks for detection on an Agilent Quadrupole Time-Of-Flight 6546 mass spectrometer. The gradient started at 85% (15% B) and decreased to 70% A over 2 minutes and then 52% A over 30 seconds. The gradient then slowly decreased to 18% A over 12.5 minutes and then 1% A in one minute and these concentrations held for four minutes. The gradient is then restored to 85% A and the column is washed for 5 minutes. Lipids were identified using MSDIAL 4.7 (PMID: 25938372) Each lipid species was quantified using the moles of corresponding class-specific lipid standards present in Splashmix.

### Short-hairpin RNA (shRNA) design and lentivirus production

shRNA against consensus coding sequence of target genes were designed using available GPP Web Portal (https://portals.broadinstitute.org/gpp/public/). The shRNA oligonucleotides consisting of sense, hairpin loop and antisense were flanked by restriction sites AgeI and EcoRI for subcloning. The shRNA oligos (ThermoFisher) were annealed and subcloned into pLKO-TRC vector between the two restriction sites. The resulting vector expressed shRNA under transcriptional control of U6 promotor along with a BFP (blue fluorescent protein) expressed under the control of hPGK promotor to ensure the delivery of plasmids to cultured neurons. The pLKO plasmid with desired oligonucleotides was delivered to HEK 293FT cells along with two packaging plasmids (pPAX2 and pMD2) using JetPRIME DNA/siRNA transfection reagent (Polyplus, 101000027). Transfection media was replaced with virus production media after 16 hours of delivery of the plasmids. The supernatant containing lentivirus was collected at 46-48 hours, filtered through 0.45 μm syringe filter set and concentrated using Lenti-X concentrator (Takara Bio, 631231). The virus titer was measured using Lenti-X GoStix Plus assay kit (Takara Bio, 631280).

### Western blot to confirm knockdown

Cortical neurons obtained from rat pups aged 0-1 days were cultured according to above procedure on 35 mm dishes coated with 50 μg/ml poly-d-lysine. At days *in-vitro* 2 (DIV-2), lentivirus was introduced to the cortical neurons. To extract proteins, neurons were scraped into RIPA lysis buffer (25 mM Tris-HCl pH 7.6, 150 mM NaCl, 1% NP-40, 1% sodium deoxycholate, and 0.1% SDS) supplemented with complete protease inhibitor (Roche, 11697498001). The protein concentration in the lysed cytosol was quantified using the BCA method, following the manufacturer’s guidelines. The proteins in the samples were denatured using sample buffer containing SDS (BioRad, 1610747) and subsequently transferred to a PVDF membrane (Millipore Sigma, IPFL00010). These membranes were then blocked with a blocking buffer (LiCor, 927-60001) for a duration of 1 hour at room temperature. After blocking, the membranes were subjected to overnight incubation with primary antibodies at 4°C, followed by HRP-conjugated secondary antibodies (Invitrogen, 31430 & 31460) for 1 hour at room temperature. Both the primary and secondary antibodies were diluted in LiCor blocking buffer according to the manufacturer’s recommendations. The HRP signal was detected using the ECL kit (ThermoFisher, 32109) and images were captured using the LiCor Odyssey Fc system.

### Fatty acid induction to hippocampal neurons

Long-chain free fatty acids, owing to their low solubility to aqueous medium, are complexed with albumin to facilitate their delivery to cells. FA-free BSA was overlayed on 0.1 M Tris (pH 8) at 37°C and it was allowed to dissolve completely. Oleic acid was added to the Tris-BSA solution in 6:1 molar ratio and stirred gently at 37°C to prepare the oleate-BSA complex. This complex was subsequently passed through a 0.22 μm syringe filter before being utilized in cell culture. Palmitic acid, in contrast to oleic acid, is solid at room temperature. FA-free BSA was dissolved in 0.9% NaCl at 40 °C, and palmitic acid was solubilized in 0.1 M NaOH by heating to 70°C. These two solutions were then combined in a 5:1 molar ratio (palmitate-BSA) and vortexed at 40°C until the solution became turbid. Similar to the oleate complex, the palmitate complex was also passed through 0.22 μm syringe filter before being used in cell culture. Oleate complex were used within 3 months after preparation, while palmitate complex was freshly prepared and maintained at 37°C during the experiment.

### BODIPY 558/568 C12 tracing in neurons

Sparsely cultured hippocampal neurons were pulsed with 10 μM BODIPY 558/568 C12 (referred as Red-C12 in present manuscript) for 24 hours in 5 μM KLH45 containing feeding media. The neurons were transferred to complete feeding media without Red-C12 for 1 hour to incorporate the Red-C12 into neuronal LDs. The neurons were subsequently washed and transferred to HBSS and chased for the defined period (0-4.5 hours) either in absence or presence of 10 μM Etomoxir or 5 μM TTX. At 30 minutes before the end of incubation time, add 50 nM MitoTracker to the media and return the neurons to incubator. The neurons were washed once with HBSS and transferred to fresh HBSS (with inhibitors) conditioned in incubator. The coverslip with the neurons were mounted on custom designed chamber for imaging on confocal microscope.

### Imaging with live neurons

Unless specified otherwise, all epifluorescence time-lapse images were acquired using custom-modified Zeiss Axiovert 200 microscope equipped with laser illumination, an Andor iXon camera (model: DU-897U-CS0-BVF) and 40X 1.3 NA Fluar Zeiss objective. To control the OBIS solid-state 488 nm and 561 nm lasers, acousto-optic tunable filters were employed on the light path. To control the buffer flow to the neurons, coverslips with cultured and transfected neurons were mounted with a laminar flow perfusion system maintained at 37°C. The perfusion solution consisted of Tyrode’s solution, comprising 119 mM NaCl, 2.5 mM KCl, 2 mM CaCl_2_, 2 mM MgCl_2_, 50 mM HEPES (pH 7.4), and 0-5 mM glucose. It was supplemented with 10 mM 6-cyano-7-nitroquinoxalibe-2, 3-dione (CNQX) and 50 mM D,L-2-amino-5-phosphonovaleric acid (APV) acquired from Sigma to suppress post-synaptic responses. For experiments involving fatty acids, glucose in the Tyrode’s solution was replaced with mentioned amount of BSA-complexed palmitate or oleate. The following concentrations of the pharmacological inhibitors were added to the appropriate Tyrode’s solution and perfused for 5-10 minutes before electrical stimulations-KLH45: 5 μM, Oligomycin: 2 μM, Etomoxir: 20 μM, Tetrodotoxin: 2 μM. Action potentials in neurons were triggered by brief 1 ms pulses, generating a field potential of ∼10 V/cm, using platinum–iridium electrodes.

### Statistical analysis

Image sequences acquired on epifluorescence microscope were analyzed using ImageJ/Fiji software embedded with plugin ‘Time Series Analyzer V3.0’ (https://imagej.nih.gov/ij/plugins/time-series.html). Approximately 20-25 regions of interest (ROIs), each covering an area of 12-18 μm^2^ and corresponding to responsive synaptic boutons, were manually selected for fluorescence measurement over time. All statistical analysis was performed with Origin 2021-23 or GraphPad Prism v6.0. Unless otherwise indicated, the data points were assumed normally distributed and were analyzed with unpaired samples t-tests. The datasets (traces and data points) are represented as mean ± SE (standard error of mean), or else indicated otherwise in the figure legends. Significance levels were represented as follows: ns for > 0.05, * for p ≤ 0.05, ** for p ≤ 0.01, *** for p ≤ 0.001, and **** for p ≤ 0.0001. The number of experimental replicates (N) used for imaging in each condition is mentioned in the corresponding figure legends.

## Acknowledgments

We thank Rina Davidson and David Simon for providing mouse brain sections used in IHC, Rachel McAllister for providing critical insights on lipidomics and members of the Ryan lab for critical discussions. This research was supported in part by NIH grants (NS036942 & NS11739) to TAR and in part by Aligning Science Across Parkinson’s ASAP-000580.

## Author Contributions

MK & TAR designed the experiments and wrote the manuscript. Neuronal cell culture experiments, cellular imaging and purification of LDs were carried out by MK. Large scale primary neuronal cultures generating LDs were carried about by DH. Electron microscopy was carried out by YL under the supervision of PDC. Lipidomic analysis was carried out by JK under the supervision of KG.

**Supplementary Figure 1.**
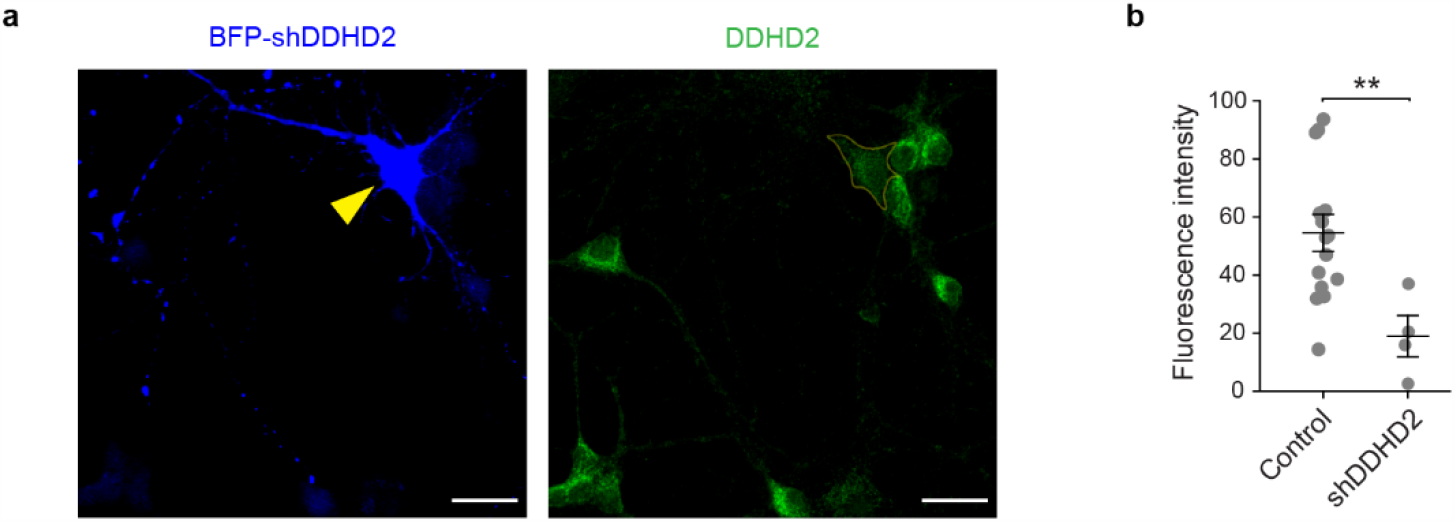
Validation of specificity of DDHD2 antibody by DDHD2 targeting shRNA in dissociated hippocampal neurons. **a**, Confocal micrographs of BFP-shDDHD2 expressing neurons immunolabeled with DDHD2 antibody. The representative shRNA expressing neuron (yellow arrowhead) shows reduced intensity of DDHD2 compared to non-expressing neurons in the same field of view. scale bar: 20 μm. **b**, Quantification of fluorescence intensities of DDHD2 in soma of control (shRNA non-expressing) and transfected (DDHD2 specific shRNA expressing) neurons. Data is represented as mean ± SE, *p*-value (** ≤ 0.01) was determined using unpaired samples t-test.

**Supplementary Figure 2.**
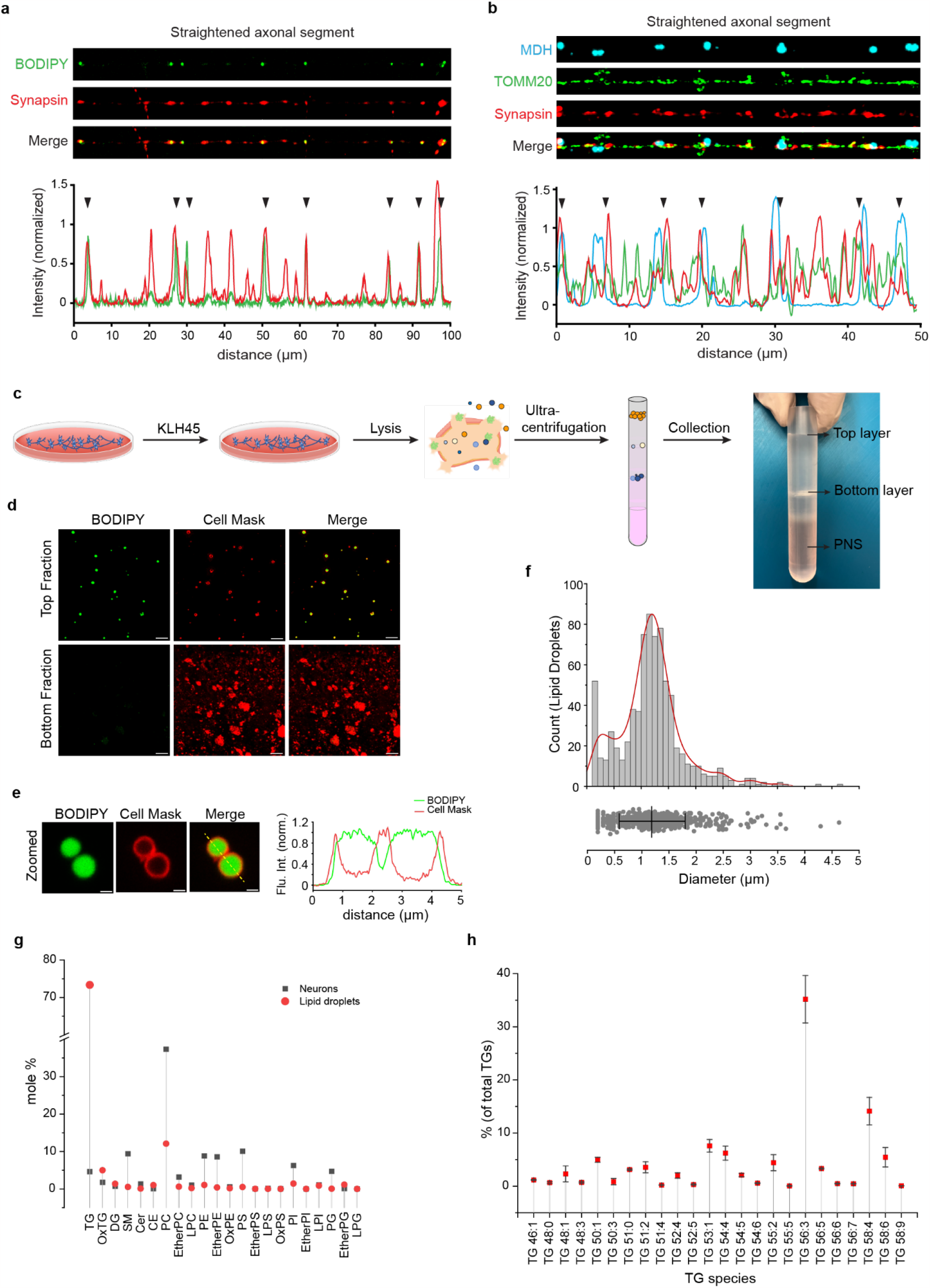
Purification and lipid profiling of KLH45 induced LDs. **a**, Confocal images of a straightened 100 μm long segment of hippocampal axon (from Fig. 2a) immunolabeled with anti-synapsin antibody and stained with BODIPY dye. Merged image and line intensity profile show repeated co-existing peaks (black arrowheads) of BODIPY (LDs, green) and synapsin (synaptic boutons, red), **b**, Confocal images of a straightened 50 μm long segment of hippocampal axon (from Fig. 2e) stained with MDH (LDs, Cyan) and immunolabeled with anti-TOMM20 (mitochondria, green) and anti-synapsin (synaptic terminal, red) antibodies. Merged image and line intensity profile show repeated peaks (black arrowheads) of MDH (cyan) adjacent to the peaks of TOMM20 and synapsin. **c**, Schematic illustration of experimental steps to induce the formation of LDs in cortical neurons and their isolation using density-based floatation in step-gradient of sucrose. The floating LDs are collected from the top layer of sucrose-free MEPS buffer (see detailed method^13^). The bottom layer contains relatively dense bi-layered membranous organelles with aqueous core. **d**, Confocal micrographs of purified LDs (top layer) and membranous organelles (bottom layer) stained with BODIPY (neutral lipid dye) and CellMask (amphipathic membrane dye). Absence of only CellMask positive, and BODIPY negative structures in top-fraction confirms the purity of LDs. Absence of detectable BODIPY-positive structures in the bottom fractions confirms that the loss of LDs was negligible during the process of LD floatation. Scale bar: 10 μm. **e**, Zoomed-in confocal micrographs and intensity profile (along yellow line) of BODIPY and CellMask stained LDs show core of neutral lipids enclosed with intact layer of phospholipids. Scale bar: 1 μm. **f**, Diameter of LDs quantified from the confocal micrographs of BODIPY stained LDs. The data is represented as mean ± SD. The mean diameter of LDs estimated is 1.2 μm. **g**, Lipid species quantified (shown as mole %) from the purified LDs compared to whole neuron after induction with KLH45. The mean values for each lipid class calculated from two independent experiments are shown. **h**, Comparative abundance (in percent of total TGs) of TG species with various lengths of fatty acid chains and saturation present in the core of LDs. Data is represented as mean ± SE.

**Supplementary Figure 3.**
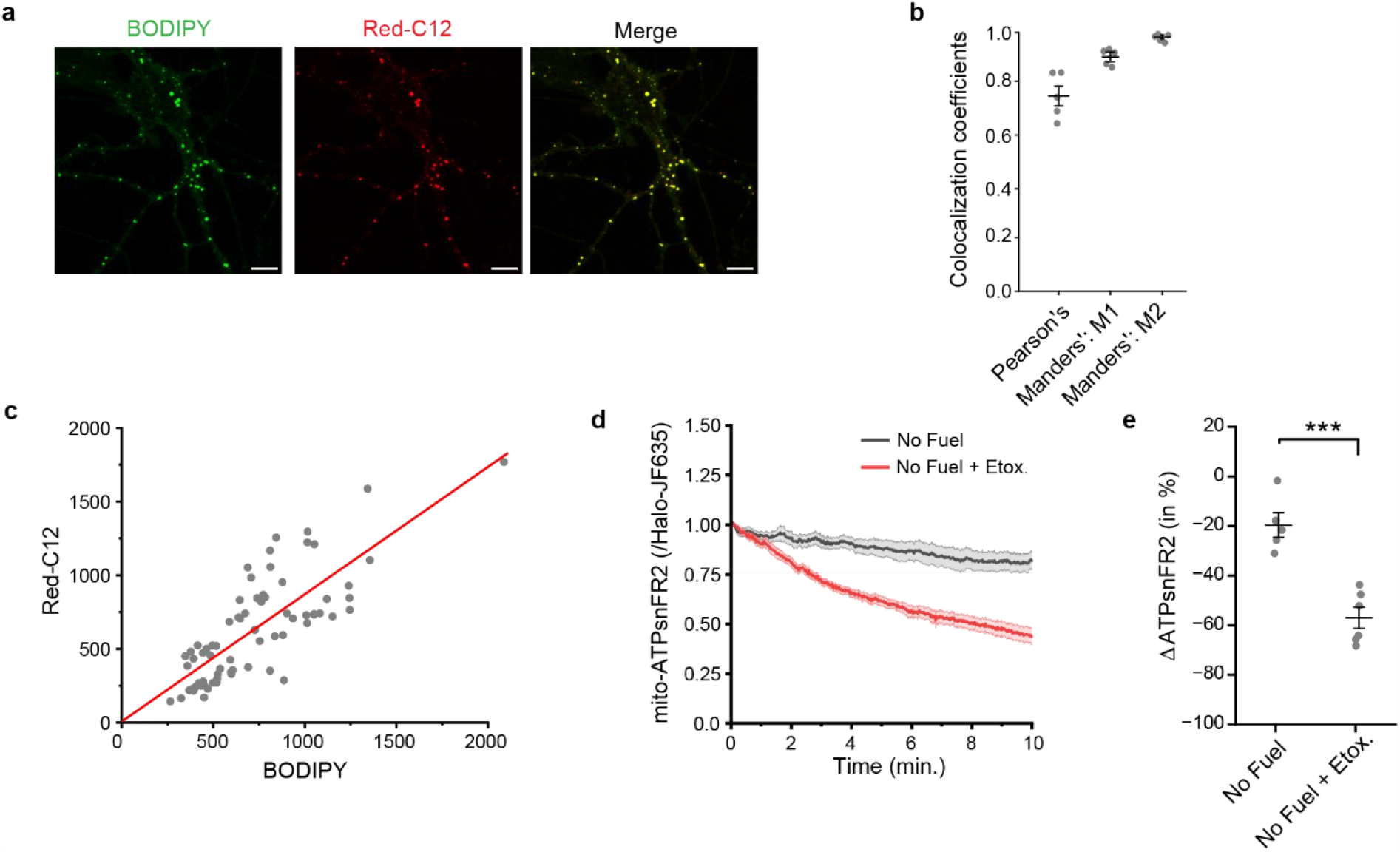
Lipid droplet derived fatty acids can help mitochondria sustain their ATP levels. **a**, Colocalization of BODIPY positive globular structures (LDs) with Red-C12 in confocal micrographs confirm the storage of exogenous fatty acids to the LDs. scale bar: 10 μm. **b**, Pearson’s colocalization coefficient quantified from the confocal images of (a). Data from N=5 is represented as mean ± SE. M1: fraction of Red-C12 overlapping with BODIPY, M2: fraction of BODIPY overlapping with Red-C12. **c**, Correlation between fluorescence intensities of BODIPY and Red-C12 measured (Pearson’s r: 0.79, R-square: 0.9) across 70 LDs. **d**, ATP levels (normalized with JF635) in the mito-ATPsnFR2-Halo expressing hippocampal neurons after induction with KLH45 followed by fuel-deprivation compared to CPT1 inhibited (by Etomoxir) conditions. Data from N ≥5 is represented as mean ± SE. **e**, Relative change in the ATP levels after 10 minutes of perfusion with the media as in (d), Data is represented as mean ± SE, N ≥5 for each condition, *p*-value (*** ≤0.001) was determined using unpaired samples t-test.

**Supplementary Figure 4.**
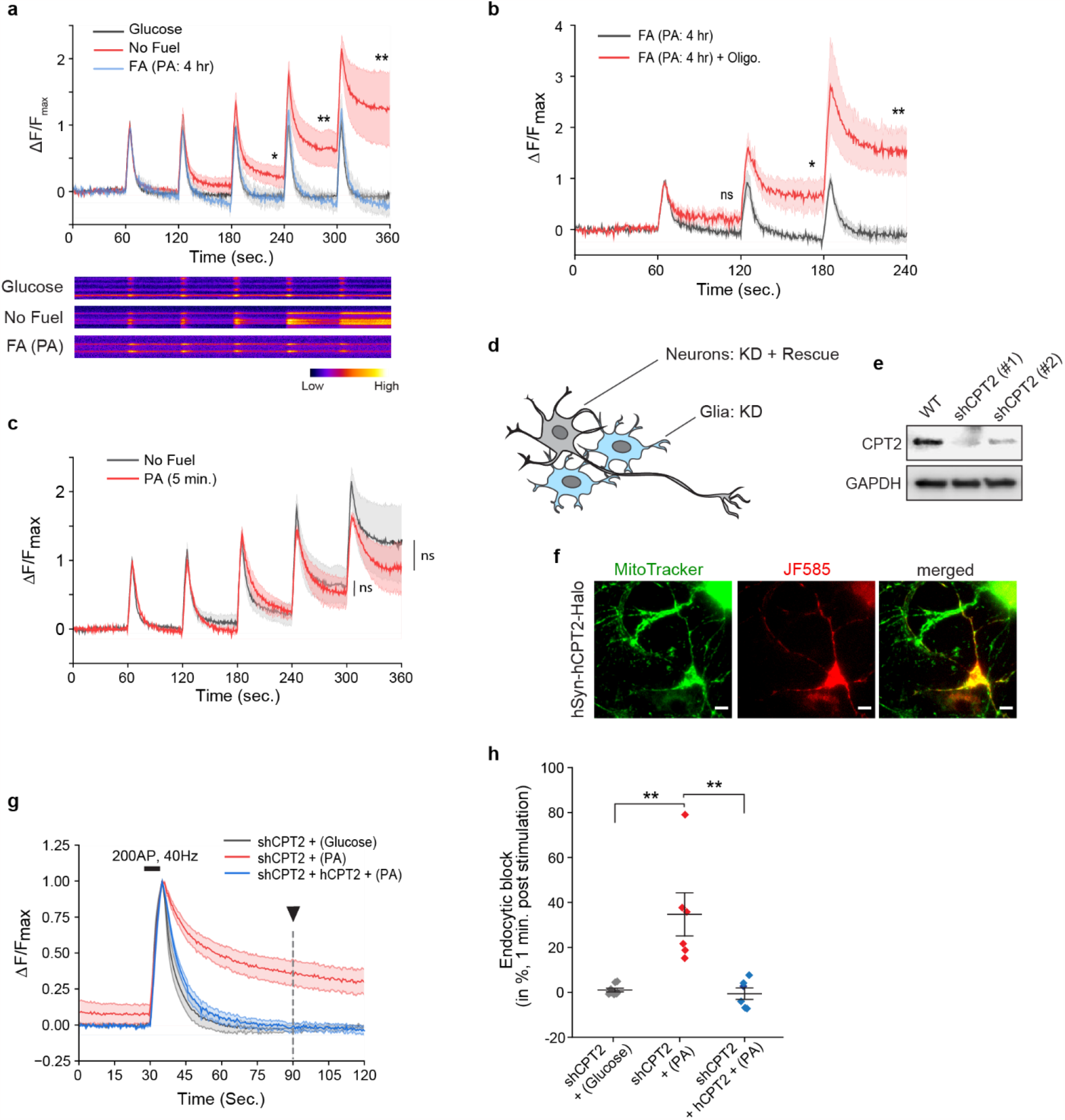
CPT1 facilitates the fueling of synaptic vesicle recycling independent of astrocytes. **a**, Fluorescence intensity (normalized with first peak) traces of vGlut1-pHluorin at synaptic boutons of the dissociated hippocampal neurons after periodic firing of action potentials (as in Fig. 4a) in absence of any external fuel, presence of glucose or pre-fed with BSA-complexed palmitate (fatty acid). Lower panels show the representative kymographs of the synaptic boutons imaged under above three conditions. **b**, Fluorescence intensity traces of vGlut-pHluorin in palmitate pre-fed or ATPase inhibited (with oligomycin) condition to palmitate fed hippocampal neurons. **c**, Fluorescence intensity traces of vGlut-pHluorin at synaptic boutons after periodic AP firing in acutely palmitate fed (5 min.) compared to no fuel condition. Traces in panels a-c represent mean ± SE. *p*-values (* ≤ 0.05, ** ≤ 0.01) were determined using unpaired samples t-test. ns signifies non-significant change in the mean values. **d**. Schematic illustration of the lenti-virus mediated knockdown of CPT2 both in glial cells and hippocampal neurons, while rescue by human synapsin promotor driven overexpression of CPT2 exclusively in neurons. **e**, Western blot to confirm the knockdown of CPT2 using two potential shRNAs. Blot of GAPDH ensures the equal loading of total proteins. **f**, Fluorescence micrographs showing colocalization of Halo-tagged human CPT2 (stained with JF585) with MitoTracker (mitochondria) in hippocampal neurons confirm the localization of the chimeric protein (CPT2-Halo) to mitochondria. scale bar: 20 μm. **g**, Normalized fluorescence intensity of vGlut1-pHluorin measured in CPT2 depleted hippocampal neurons show reduced endocytosis in palmitate fed (red) compared to glucose fed condition (grey). Expression of human CPT2 (shRNA resistant) rescues the endocytosis. **h**, Endocytic block (in %) measured after 60 seconds of AP firing (shown by black arrowhead in g). Data is represented as mean ± SE. *p*-value (** ≤ 0.01) was determined using unpaired samples t-test.

